# Blood flow directionality establishes the SCN as source and OVLT as target within a new vascular portal pathway

**DOI:** 10.1101/2023.06.20.545729

**Authors:** Ranjan K. Roy, Yifan Yao, Isabella K. Green, Rae Silver, Javier E. Stern

**Author notes:** <u>Corresponding author</u>: Javier E. Stern, M.D. Ph.D., Neuroscience Institute, Georgia State University, Atlanta, GA 30302-5030 United States.

## Abstract

The suprachiasmatic nucleus (SCN) is the locus of a brain clock that sets the phase of oscillation in cells throughout the brain and body. Anatomical evidence reveals a portal system linking the SCN and the OVLT (here termed SCN-OVLTp). This discovery begs the question of the direction of blood flow and the nature of diffusible signals that flow in this specialized vasculature. Here we show unequivocally that the direction of blood flow is from the SCN to the OVLT, that the rate of flow is under circadian regulation, and that vasopressin (AVP) is present in portal vessels following systemic injection. These findings highlight a previously unknown CNS communication pathway. It is well established that the SCN is required for circadian regulation of AVP in the CSF and that the OVLT bears AVP receptors. Specifically, SCN neurons are necessary for time-stamped signals such as the peptide AVP, that can travel via portal veins to a target in the OVLT. The OVLT, a circumventricular organ offering a “window to the brain,” can relay neural and diffusible signals to broad brain areas via its efferent connections and via the CSF. We conclude that the SCN-OVLTp, like that of the pituitary portal system, discovered almost a century ago, allows neurosecretions to reach nearby specialized target sites, thereby avoiding dilution in the systemic blood. In both of these brain portal pathways, the target site, namely the pituitary and OVLT respectively, relay signals broadly, to both the brain and the rest of the body.

## INTRODUCTION

The portal pathway linking the capillary beds of the hypothalamus and pituitary was first described in 1933 and was long believed to be the only such vascular pathway in the brain(1). Almost 90 years later, that unique position changed with our discovery of a second portal pathway connecting the mouse SCN and the organum vasculosum of the lamina terminalis (OVLT), a nearby circumventricular organ (CVO)(2). These findings point to the mechanism whereby small populations of neurons can produce locally effective concentrations of secretions that reach their targets via this specialized vascular system, avoiding dilution in the general circulatory system. The functional significance of this new neurovascular OVLT portal pathway (here termed SCN-OVLTp) requires unveiling the direction of blood flow between these nuclei. This in turn will permit determination of the source and the target of signals traveling in this system and its functions. It is worth noting that in the case of the hypothalamic-pituitary portal system, this nexus of studies regarding the direction of blood flow took about two decades (3), and many years thereafter led to the 1977 Nobel prize in physiology and medicine to Guillemin and Schally.

A great deal has been written about both the SCN, the OVLT and CVO’s. The SCN, locus of the brain’s circadian clock, lies at the base of the brain, just above the optic chiasm, adjacent to the third ventricle. Neuronal efferents from the SCN reach numerous central nervous system (CNS) targets(4–6) including the OVLT(7). However, it is also well-established that the SCN produces neurosecretions that exhibit daily rhythms (8–10), and that diffusible signals are sufficient to sustain circadian rhythms. Thus, when transplanted into the third ventricle, the grafted SCN, even when placed within a capsule that prevents fiber outgrowth (11) can restore locomotor, drinking and gnawing rhythms in animals in which this nucleus has been ablated (reviewed in(12–14)). Additionally, in an *ex-vivo* slice preparation, SCN neuropeptides including vasoactive intestinal peptide (VIP), arginine vasopressin (AVP) and gastrin releasing peptide (GRP) secreted by a “donor” SCN can support rhythmicity in recipient arrhythmic SCN “host” tissue (15).

Like the SCN, the OVLT lies immediately above the optic chiasm near the wall of the third ventricle. The OVLT is a sensory CVO bearing fenestrated capillaries, and is enriched in receptors for hormones and neuropeptides (16, 17). The fenestrated capillaries allow systemic circulating factors to permeate the nucleus and to reach the sensory neurons of the OVLT. Information can then be relayed from OVLT to numerous targets, including the supraoptic nucleus (SON), paraventricular nucleus (PVN), median preoptic nucleus (MnPO) and the SCN itself (17, 18) via its neural efferents, and to the CSF via its leaky blood vessels. The OVLT is implicated in a variety of centrally regulated processes many of which have daily fluctuations including, anticipatory locomotor activity, anticipatory thirst and hunger, ovulation, osmoregulation, various gonadal functions, fever and sickness behaviors (17–20).

While the route(s) travelled by diffusible signals from the SCN remains unknown, the newly discovered SCN-OVLTp stands as a potential candidate. We previously hypothesized that the flow of information is from SCN to OVLT(2). If so, then neurons of this small nucleus could secrete sufficient amounts of biologically significant neuropeptides that travel to specialized nearby targets in the OVLT. The sensory neurons and fenestrated leaky blood vessels of this CVO could then relay circadian signals broadly via their neural and vascular outputs, thereby orchestrating circadian rhythms throughout the body.

While the foregoing hypotheses are plausible, at present, we do not know whether the SCN-OVLTp occurs in animals other than mouse, and whether the direction of blood flow is from the SCN to the OVLT or vice versa. Here we assessed the former question by studying the rat, and we next established the direction of blood flow to assess the source and target within the SCN-OVLTp system. We then explored, in both sexes, circadian regulation of blood flow in the portal vessels and finally, we asked whether the SCN neuropeptide AVP could enter portal vessels following systemic administration.

## Results

### Anatomy of the rat SCN-OVLT portal pathway

To determine whether there exists a portal pathway between the SCN and OVLT in rat, we visualized the SCN and the vasculature of the ventral aspect of the brain using iDISCO cleared material, immunochemistry and light sheet microscopy, as previously reported for the mouse(2) (see methods pipeline in ***Supplementary Fig.1***). ***Fig.1*** shows the major features of the volume between the two structures of interest. The scan in the left panel (***Fig.1a*** provides a 3D view of the location of the SCN and OVLT within the third ventricle. Next, the scans of three separate channels and their merged images (***Fig.1b*** identify the SCN by its characteristic AVP neurons, the vasculature of the entire region labelled with collagen, and the arterial vasculature marked by smooth muscle actin (SMA) staining and a merged image of the three labels. The final panel shows the traced blood vessels that connect the SCN and OVLT. The results indicate that the major features of the rat portal pathway are similar to that of the mouse, with fine portal capillary vessels coursing along the floor of the third ventricle between the SCN and OVLT.

**FIGURE 1.**
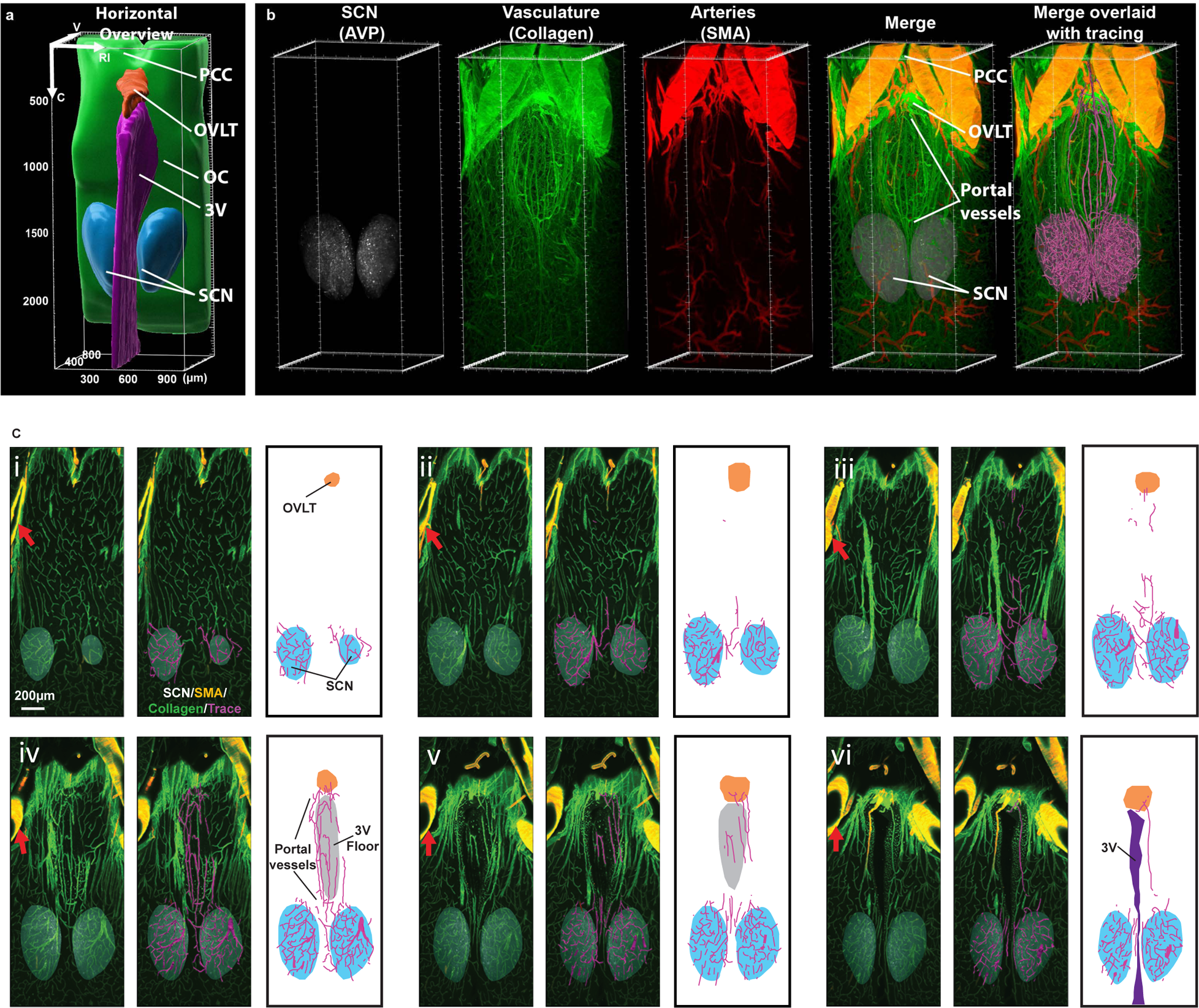
Anatomy of the rat SCN-OVLT portal system. **a,** The image shows the orientation of the horizontal scan that was done on the tissue shown in **b**. In the rostrocaudal axis, the scanned volume extends from the prechiasmatic cistern (PCC) to retrochiasmatic area (Bregma −0.84 mm to 1.72 mm)(34) and includes both sides of the SCN and its entire ventral-to-dorsal extent. Reference axis denotes the orientation of the tissue; C=caudal; V=ventral; D=dorsal; RI=right. OVLT = orange, SCN = blue, 3V= magenta; OC = green**. b**, The image shows immunostaining and reconstruction of portal vessels connecting SCN and OVLT. The near-horizontal axis is for the same tissue depicted in A. Panels from left to right show the SCN identified by AVP staining, the full vasculature of the region labelled with collagen, arteries labelled with SMA, the merged image for the three markers and last, the computer assisted tracings of blood vessels merged with all prior panels. **c**, The figure demonstrates the portal capillaries from their most ventral to most dorsal aspect in serial 50 µm optical slices. The plates show triplets of images as follows: Left panel=merged AVP (white), collagen (green) and SMA (yellow); Middle panel=blood vessel traces (magenta) are superimposed on immunochemical results of the left panel; Right panel=drawing identifying structures shown in the middle panel. Details of the serial plates are as follows: (i) Initial appearance of the OVLT and portal blood vessels in the ventral SCN. (ii –iii) Portal vessels of the SCN extend rostrally from the SCN. (iv-vi) The rostrally running vessels of the SCN travel along the floor of the 3V and join the OVLT. Portal vessels do not occur in the more dorsal aspect of the SCN (not shown). For orientation in the serial sections, the red arrows in the left panel of each triplet points to a blood vessel that is seen in all the photomicrographs demonstrating the continuity of the optical slices. 3V=purple, AVP=white, collagen=green, SMA= red, overlay of red/green=yellow, traces=magenta. Abbreviations: AVP=Arginine vasopressin, OC=optic chiasm; OVLT=vascular organ of lamina terminalis; PCC=prechiasmatic cistern; SCN=suprachiasmatic nucleus; SMA=smooth muscle actin, 3V=third ventricle. See also Figure S1.

To demonstrate the precise course of the SCN-OVLTp vessels as they travel between the SCN and OVLT, we prepared serial horizontal optical scans through the dorso-ventral extent of the entire region (***Fig.1c***). Triplets of images are shown from the ventral-most aspect of the SCN at which portal vessels are observed through to the dorsal-most aspect of the SCN bearing these vessels. There are no portal vessels connecting to the SCN in the optical scan below and above these regions (*not shown*).

Both nuclei of interest have complicated shapes that change through their volumes in each axis. To guide the placement of the microscope for in vivo imaging, we performed precise measurements of the depth of the SCN and OVLT above the optic chiasm, the distances between these structures, and the cross-sectional size of the portal vessels (***Fig. 2***). In horizontal view the portal vessels are readily seen close to the midline of the brain (**Fig.2a**). In sagittal view the distance between the SCN and OVLT is short, and both nuclei can be identified and visualized in the same scan (***Fig.2b***), obviating the need to register the scan against a template. The differing relative depth of the SCN and OVLT above the optic chiasm is also evident. Detailed measurements of these structures is provided in the legend of ***Fig. 2***.

**FIGURE 2-.**
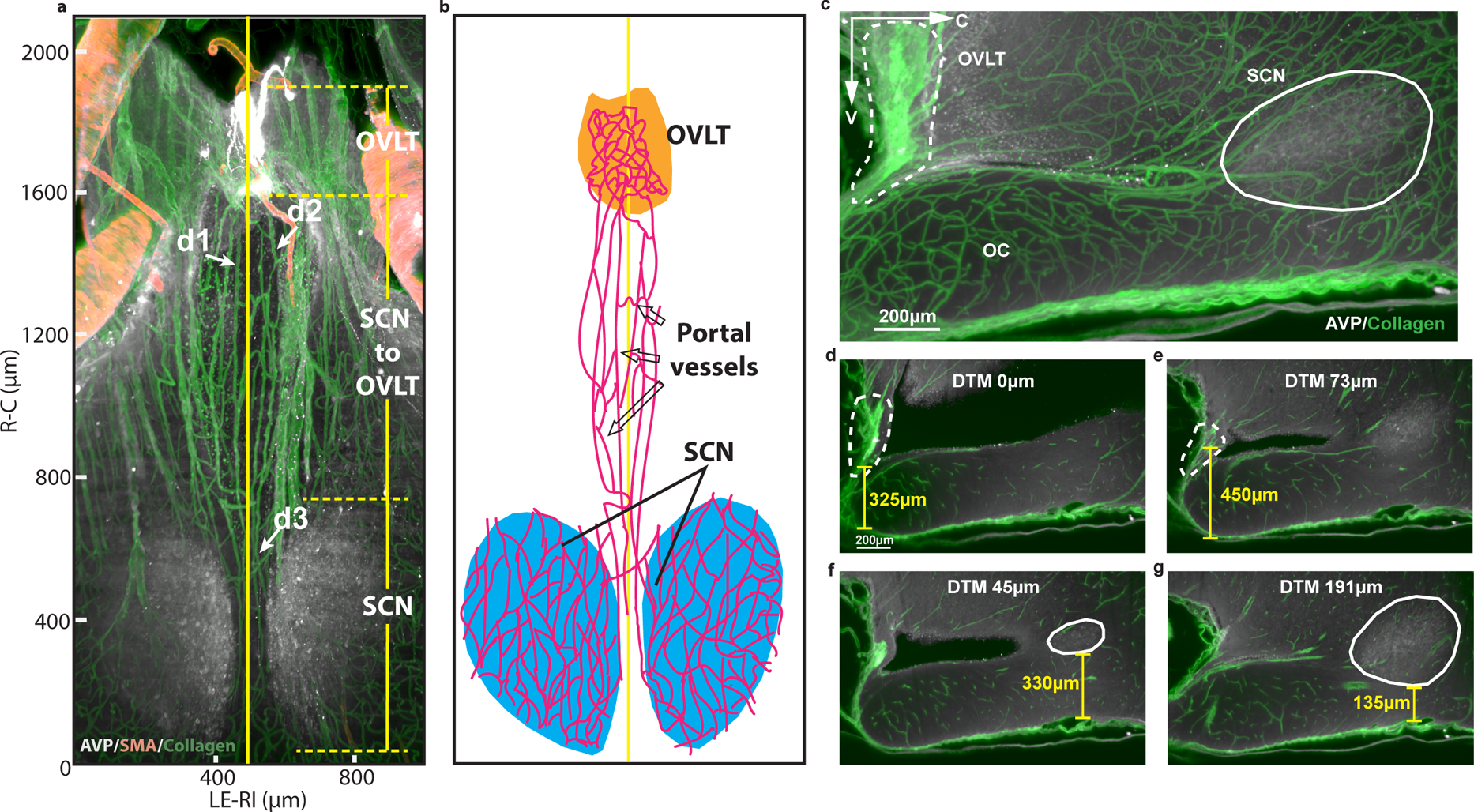
Size and locations of the SCN, OVLT and portal vessels. **a,** Vasculature of the SCN and OVLT region in a 100µm optical horizontal slice. The rostrocaudal (R-C) extent of OVLT is 270µm; the distance between caudal OVLT and rostral SCN is 730µm; the R-C extent of SCN is 640µm. The vertical yellow line at 490µm on the left-right (LE-RI) axis indicates the mid-line of the brain. Portal vessels lie within 100µm of the midline. The diameter of portal vessels at d1,2,3 are as follows: d1 = 11µm, d2 = 13µm, d3 = 8µm. **b**, The drawing identifies structures shown in A, and highlights the portal vessels near the midline (yellow vertical line) of the brain. **c** Vasculature of sagittal optical slice (150µm) of the SCN-OVLT region. To measure the height over the OC of the OVLT (dashed line) and SCN (solid line) 2µm thick slices were captured. **d**, The shortest distance from OVLT to the ventral OC is 325µm and occurs at the midline (distance to midline [DTM] =0)**. e,** The greatest distance from ventral OVLT to ventral OC is 450µm, and at this point, the DTM is 73µm. **f**, The greatest distance between ventral SCN and ventral OC is 330µm, and here the DTM is 45µm. **g**, The shortest distance between ventral SCN and the ventral OC is 135µm, and here the DTM is 191µm.

### In vivo 2-photon imaging of the eGFP-AVP neurons in the SCN and the SON

We identified the SCN based on the presence of dense endogenous eGFP-AVP fluorescent neurons and fibers, using a modified version of our surgical approach to expose the ventral hypothalamus for in vivo imaging(21). As shown in ***Fig. 3a***, the SCN was visualized medially to the SON and contains eGFP-AVP fluorescent neurons of much smaller size than those observed in the SON. In a few cases (n=3) both SCNs and SONs were exposed bilaterally (***Supplementary Fig.2a***). Based on the anatomical characteristics of the portal system described above, we focused our *in vivo* imaging studies on the rostral aspect of the SCN.

**FIGURE 3.**
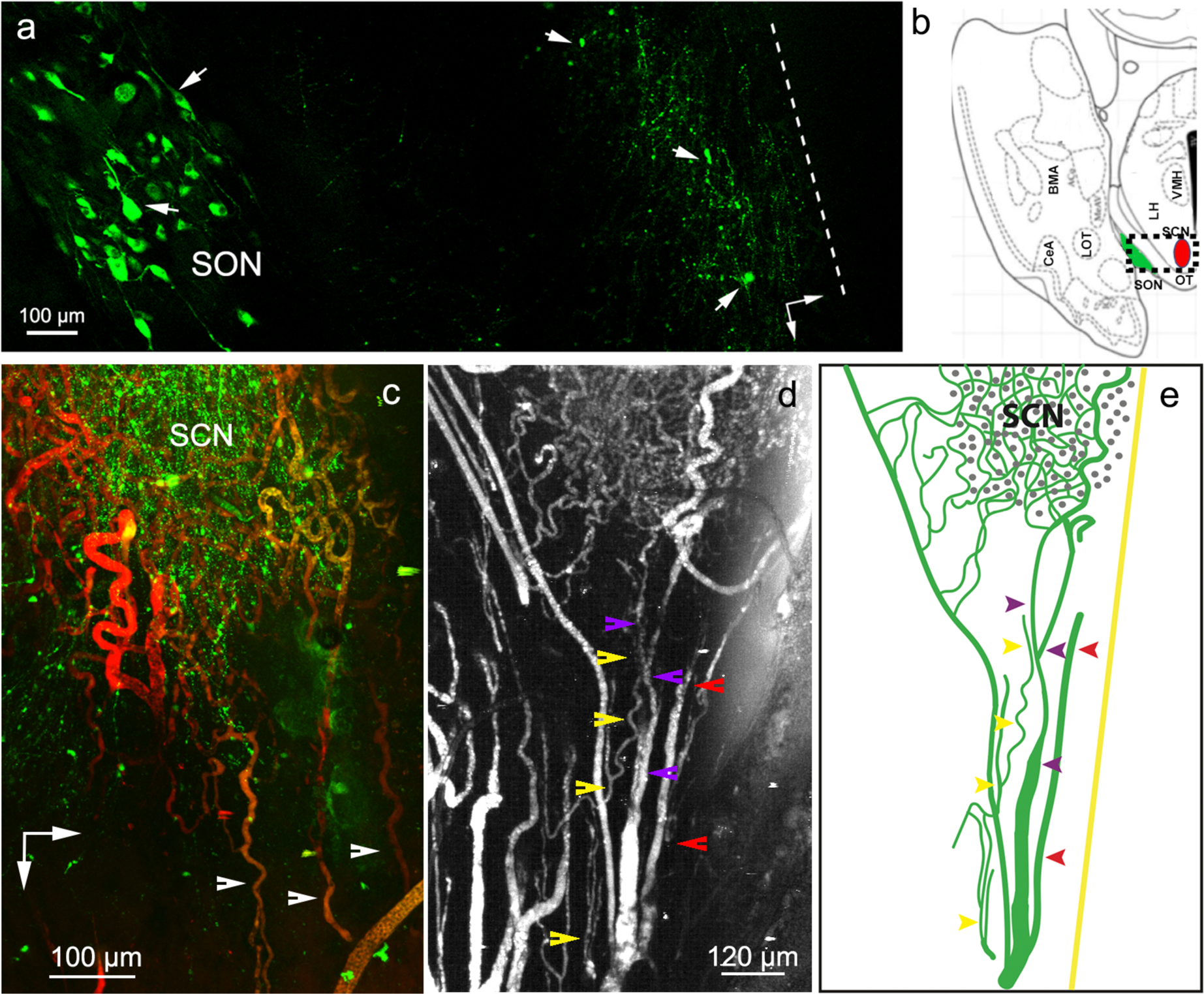
*In vivo* 2-photon imaging of the SCN microvasculature. **a**, Composite of sequential medio-lateral images (16X objective) showing eGFP-AVP neurons and fibers in the right SON and SCN of an eGFP-AVP rat (arrows). Note the large magnocellular neurons in the SON. Dashed line represents the apparent midline. Vertical and horizontal arrows point rostrally and medially, respectively. **b**, Cartoon of the brain atlas(34) showing a horizontal section of the brain and the area imaged (square). The SON and SCN are highlighted in green and red, respectively. **c**, *In vivo* 2-photon imaging of the rostral aspect of the SCN following rhodamine 70 kDa dextran infusion (I.V.). Note the dense capillary network within the SCN (containing dense endogenous eGFP-VP fluorescence) and the horizontal, rostro-caudally running vessels (arrowheads). Vertical and horizontal arrows point rostrally and medially, respectively**. d**, Representative FITC 70 kDa dextran-l<u>abelled</u> capillary vessels of the SCN and the rostro-caudally running portal vessels lying close to the midline (arrowheads; each color indicates a separate vessel). The image is shown in black and white to enhance visualization. **e**, drawing identifying the key structures shown in the middle panel b. The yellow straight line represents the location of the brain midline. See also Figure S2.

### *In vivo* 2-photon imaging of the SCN microvasculature

Vascular filling with rhodamine 70 kDa dextran I.V. revealed a dense capillary network at this rostral region of the SCN, with only a few larger vessels noted (***Fig.3c*)**. In agreement with the anatomical results, a distinct set of blood vessels coursed very close to the midline between the rostral aspect of the SCN, and the caudal aspect of the OVLT (***Fig.3d***). A different example at lower magnification is shown in ***Supplementary Fig.2b***. The mean diameter of these vessels was 11.4. ± 0.6 µm (n=30).

In confirmation that these were portal venules and not arterioles, rats were first given an I.V. injection of the artery-specific dye Alexa Fluor 633(22) to pre-label parenchymal arterioles (***Fig.4a,d***). Next, vascular filling was examined after I.V. administration of either rhodamine 70 kDA (see examples shown in ***Fig.4b,c***) or FITC 70 kDA dextrans (see examples shown in ***Fig.4d,e***) to label all the microvasculature (see additional samples in ***Supplementary Fig 2c-f***). Using this approach, we found only a few Alexa Fluor 633-labelled arterioles within the rostral SCN, and in all cases (n=8), the rostrally running portal vessels were negative for Alexa Fluor 633. In summary, these properties of the *in vivo* imaged material, and their correspondence to the anatomical features and measurements shown in ***Fig. 1,2***, confirm the identity of the SCN-OVLTp venules *in vivo*.

**FIGURE 4-.**
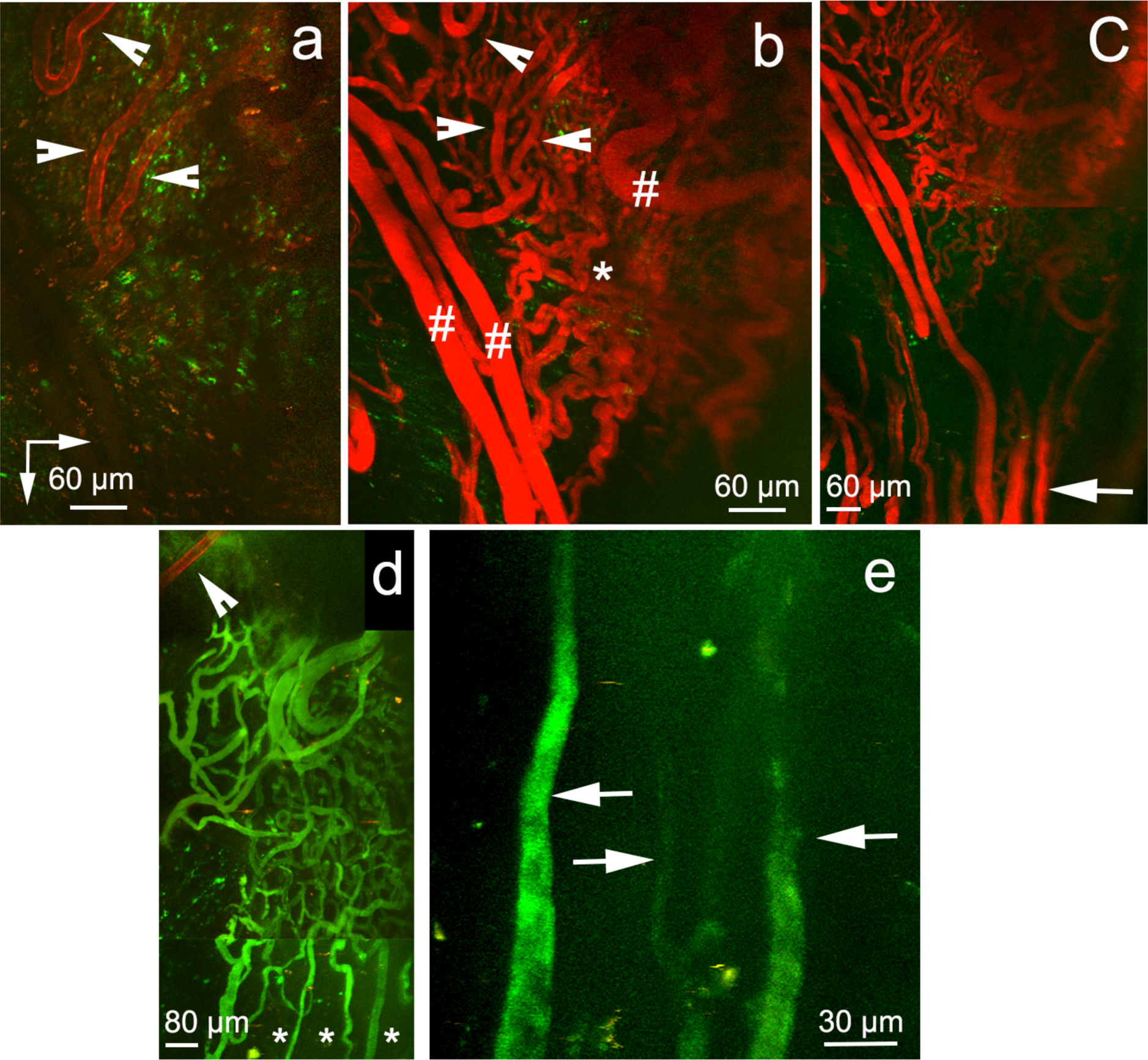
The SCN-OVLT portal vessels are venules and not arterioles. **a,** 2-photon image of the rostral aspect of the SCN after an I.V. injection of Alexa Fluor 633. Few arterioles show positive staining (red, arrowheads). **b**, Same region shown in **a**, shown at a lower magnification following labeling of the entire microcirculation with rhodamine 70 kDa dextran infusion (I.V.). Arrowheads point to the arterioles shown in **a**. The asterisk and hashtag indicate capillaries and venules, respectively, within the SCN. **c**, Composite of images at lower magnification (4X objective) of the same region shown in **a** and **b**, to show the bundle of portal vessels labelled with rhodamine 70 kDa dextran (arrow). Note the absence of Alexa Fluor 633 staining in the SCN capillary network or any other vessels running rostrally from the SCN. **d**, 2-photon image composite of the rostral SCN showing an Alexa Fluor 633 stained vessel (arrowhead) and vessels subsequently stained with FITC 70 kDa dextran (I.V.). Note the scarce Alexa Fluor 633 staining at this SCN level (arrowhead) and that the rest of the stained vasculature shows only FITC 70 kDa dextran staining. Asterisks indicate initial segment of a few portal vessels. **e**, Image taken more rostrally than **d**, and shown at higher magnification, to better show the portal vessels stained with FITC kDa dextran but lacking Alexa Fluor 633 staining (arrows). Vertical and horizontal arrows in a point rostrally and medially, respectively. See also Figure S2.

### Direction of blood flow is from SCN to OVLT

To determine the directionality of blood flow within the SCN-OVLTp, we performed fast sequential 2-photon imaging of the acute loading phase of the portal vessels during I.V. infusion of Rhodamine 70 kDA dextran. This allows measurement of blood flow and directionality as the vessels are stained in real time with the intravascular dye. As shown in the two samples in ***Fig. 5***, real time measurement of intravascular loading of the SCN-OVLTp venules with the fluorescent dye showed that vessel staining started at caudal segments near the SCN, and then rapidly moved rostrally toward the OVLT. A kymograph plot (Rhodamine 70 kDA dextran intensity as a function of time and distance along the portal vessels) confirmed the direction of dye movement from caudal (SCN) to rostral (OVLT) aspects. Similar results were observed in n=6 independent experiments in separate rats. Thus, real time measurement of the acute phase of fluorescent loading of the SCN-OVLTp vessels supports flow from the SCN towards the OVLT.

**FIGURE 5-.**
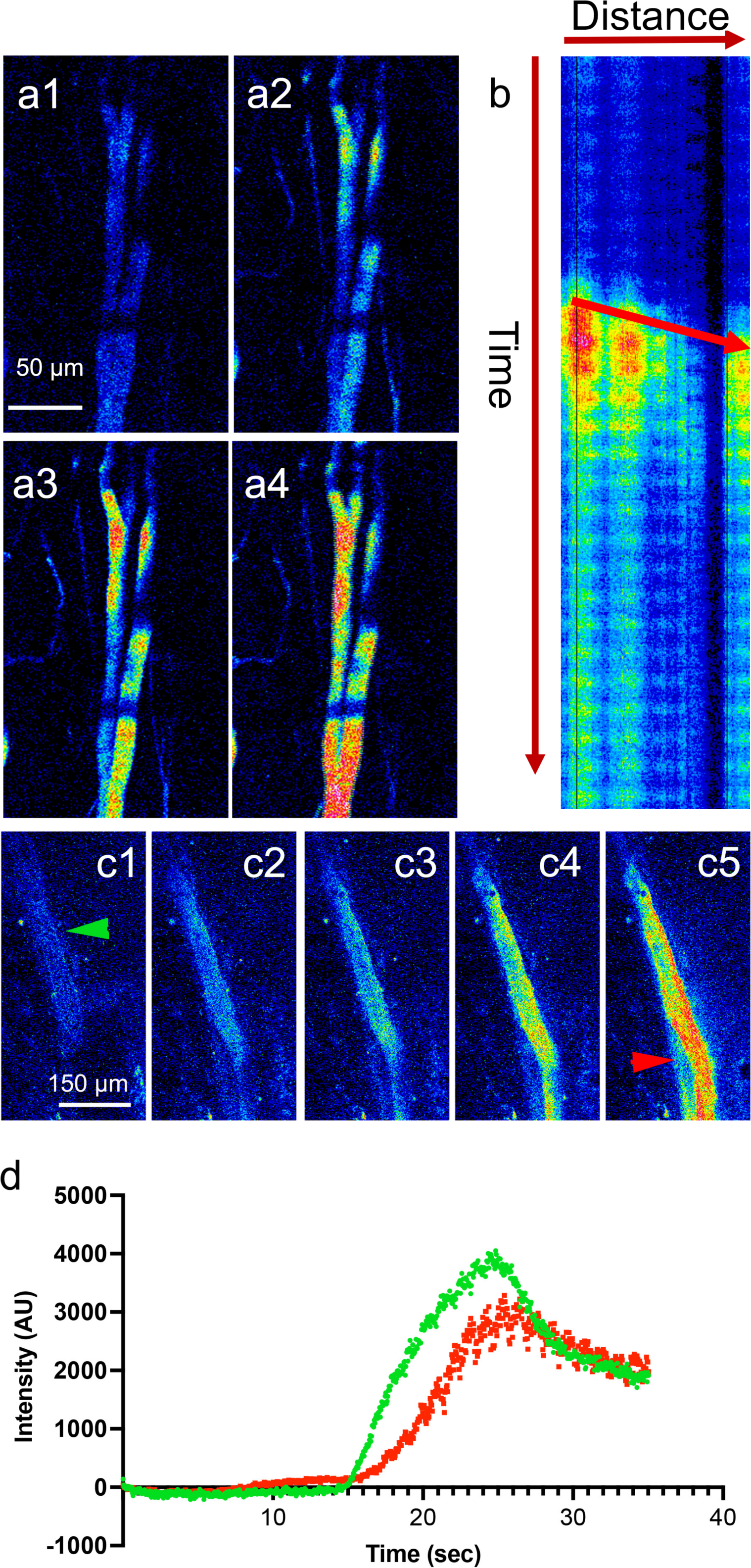
Acute loading of the SCN-OVLT portal system supports blood flowing from SCN towards the OVLT. **a,** Sequential time-series 2-photon imaging (a1-a4) of the acute loading phase of the portal vessels during rhodamine 70 kDa dextran I.V. infusion (pseudocolor). Note that loading starts at caudal segments, moving then rostrally. **b**, Kymograph plot of rhodamine 70 kDa dextran intensity as a function of time (vertical arrow) and distance (horizontal arrow, from caudal to rostral) along the portal vessels. The arrow within the graph indicates direction of movement over time, supporting a flow from the SCN (caudal) to the OVLT (rostral). **c**, Sequential time-series 2-photon imaging of a different portal vessel during the acute loading phase (rhodamine 70 kDa dextran I.V. infusion). Green and red arrows indicate segments where plots in **d** were generated. **d**, Plot of portal vessel fluorescence intensity over time taken at a caudal (green) and rostral (red) segments from the vessel shown in **c**. Note the delayed increased fluorescence in the rostral segment, supporting blood flow in a caudal-to-rostral direction, from the SCN towards the OVLT. Vertical and horizontal arrows in **a1** point rostrally and medially, respectively.

In a second complimentary approach, we measured blood flow direction and velocity by monitoring the movement of red blood cells (RBCs), as previously described(21). These appeared as moving dark spots in fluorescently labeled vessels in a segment of an imaged portal vessel. Representative images (***Fig.6a-c***) and a video (***Supplementary video 1***) compellingly show unidirectional movement of RBCs within the SCN-OVLTp system, running from a caudal to a rostral orientation. This property was observed in every rat assessed (n= 10 females and n= 13 males). The mean basal blood flow velocity in the SCN-OVLTp was 182.5 ± 19.9 µm/ms (n=16). There was no significant difference between the sexes in either velocity or diameter (p> 0.54 (t= 0.62) and p> 0.35 (t= 0.97), respectively, unpaired t test, n= 6 females and 10 males), and there was no significant correlation between the velocity of blood flow and the portal vessel diameter (***Fig.6e***), r^2^=0.007, p=0.75, n=16). An opposite direction of blood flow, from rostral to caudal, was observed in neighboring lateral lying vessels (*not shown*).

**FIGURE 6-.**
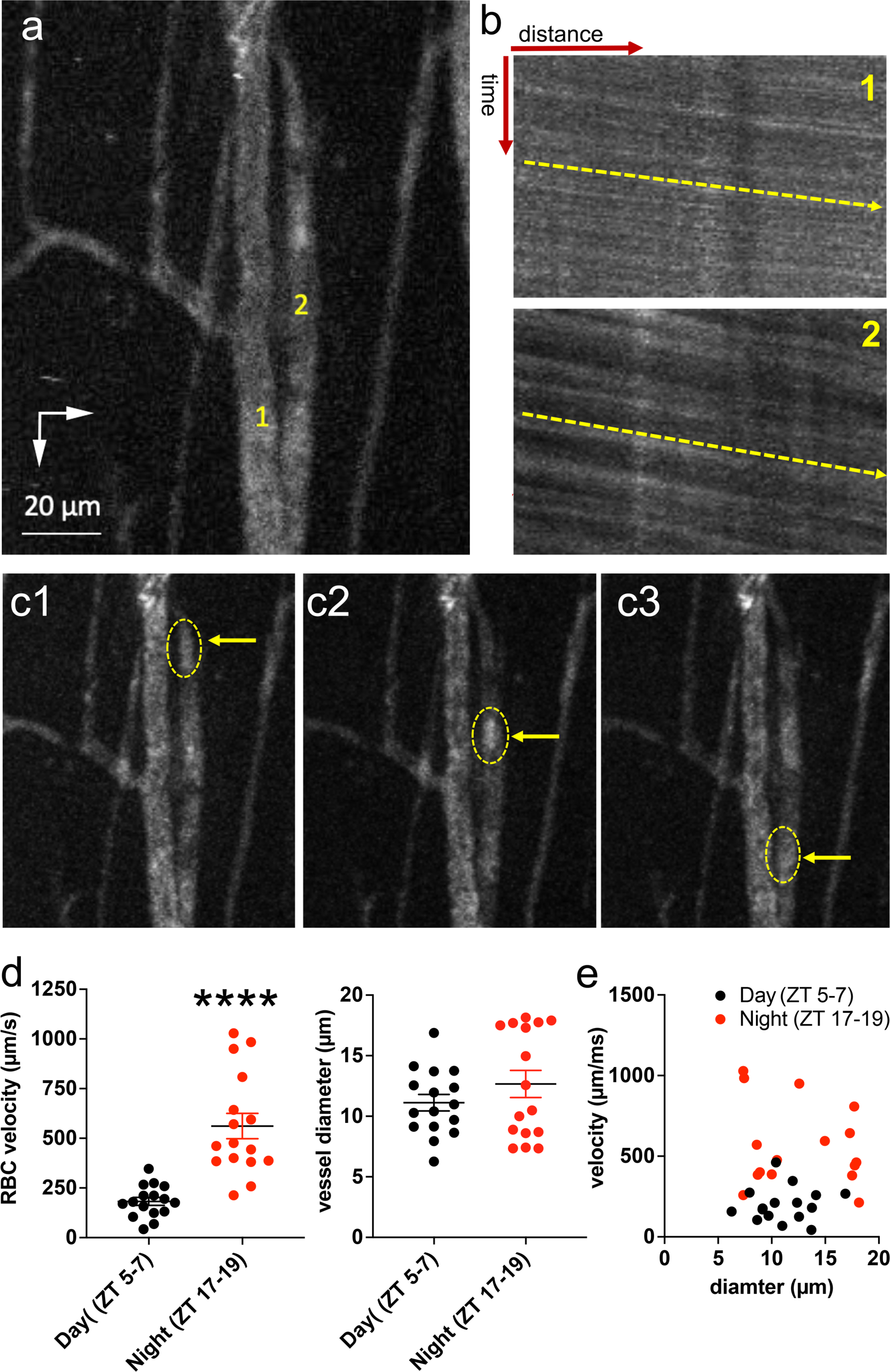
Measurements of red blood cell (RBC) movement within the SCN-OVLT portal system supports blood flowing from SCN towards the OVLT and increased blood flow during the night phase. **a,** 2-photon imaging of a pair of SCN-OVLT portal vessels (1,2) used to measure RBC movement. **b**, Kymograph plots of Rho70 kDa intensity as a function of time (vertical) and distance (horizontal, caudal to rostral) along the portal vessels 1 and 2 from A. Note the streak slopes (yellow arrows) showing movement from caudal-to-rostral over time. **c**, Sequential time-series 2-photon imaging (**c1-c3**) of the portal vessels shown in **a**, showing a group of RBCs (oval, yellow arrow) moving progressively in a caudal-to-rostral direction. **d**, Summary of RBC velocities (left) and portal vessel diameters in SCN-OVLT portal system during the day (ZT 5-7, black) and night (ZT 17-19, red) phases. **e**, Plots of RBC velocities as a function of vessel diameter at day and night phases, showing a lack of significant correlation in both cases (r^2^: 0.007 p= 0.75 and r^2^: 0.04 p= 0.46, respectively, Chi-square tests). Vertical and horizontal arrows in **a1** point rostrally and medially, respectively. ****p< 0.0001 vs day (unpaired t test). Error bars represent ±SEM.

### Time of day impacts blood flow in the SCN-OVLT portal system

To determine if blood flow within the SCN-OVLT portal system varies according to the time of day, we compared blood flow at ∼mid-day (zeitgeber time, ZT 5-7) and ∼mid-night (ZT 17-19). Remarkably, RBC velocity was significantly faster at night than in the day (ZT 17-19 vs ZT 5-7; unpaired t test, p< 0.0001, t= 5.66 ***Fig.6d***). Conversely, there were no time of day differences in portal vessel diameter between these time points (p=0.24, t= 1.18 ***Fig.6d***). Furthermore, as in the mid-day measurements, there was no significant correlation between portal vessel blood flow velocity and diameter at the mid-night time points (***Fig.6e***, r^2^=0.04, p=0.46).

### I.V. infused AVP travels within the SCN-OVLT portal system

To determine if a functionally relevant neuropeptide can penetrate and circulate within the SCN-OVLTp system, we systemically infused a fluorescently tagged arginine vasopressin (AVP_FL_). An I.V. infusion of AVP_FL_ resulted in a rapid increase in fluorescent signal within the SCN-OVLTp vessels (n=7, ***Fig.7a*** and ***Supplementary Movie 3***). Neither vessel diameter nor velocity were significantly changed by the infusion of AVP_FL_ (p> 0.78 (t= 0.29) and p> 0.20 (t= 1.88) respectively, paired t-test, ***Fig.7d***).

**FIGURE 7-.**
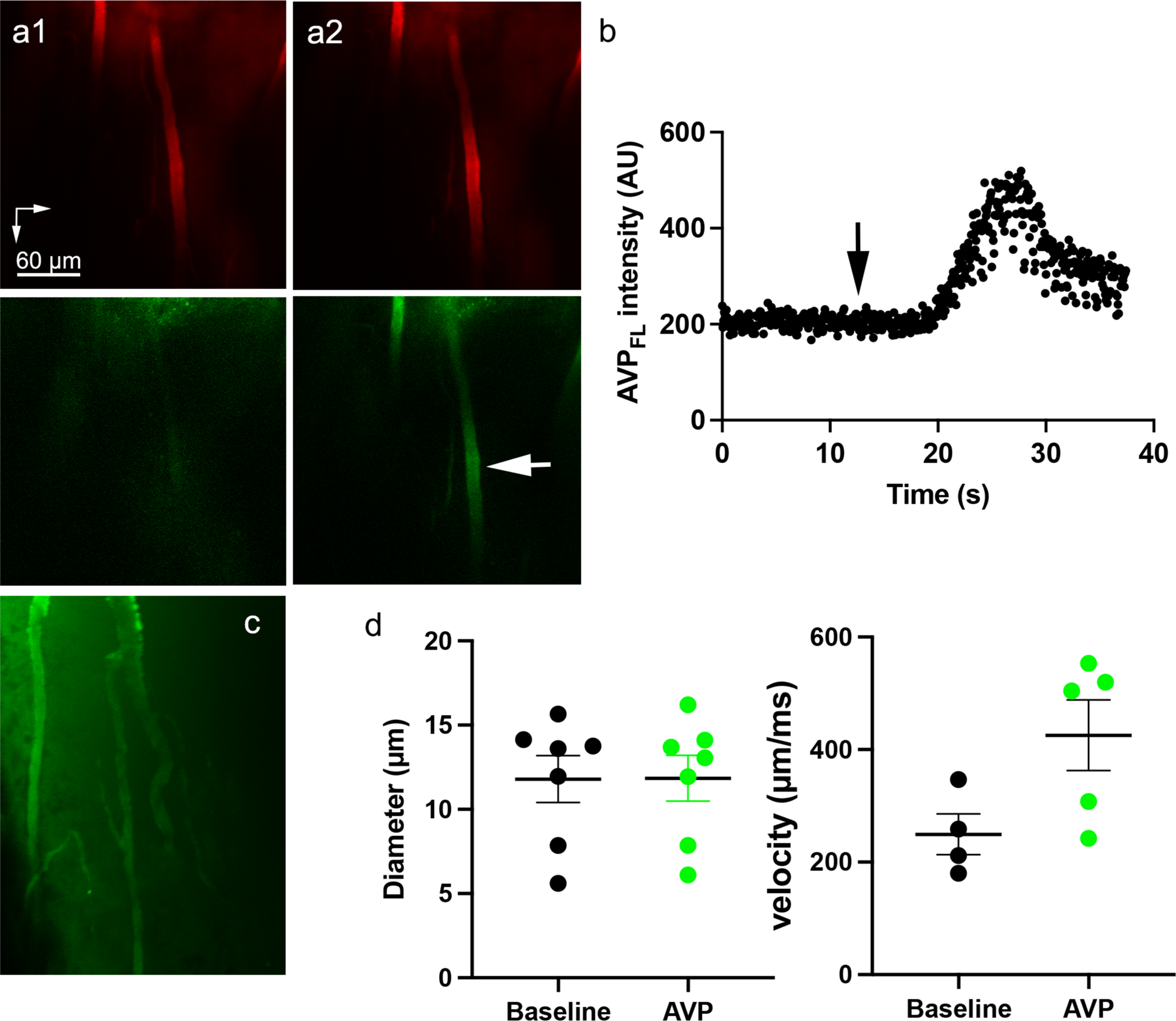
A fluorescent form of AVP (AVP_FL_) infused into the circulation can enter and travel within the SCN-OVLT portal system. **a,** 2-photon images of SCN-OVLT portal vessels pre-loaded with rhodamine 70 kDa dextran I.V. infusion (red, upper panels) before (a1) and after (a2) systemic infusion of AVP_FL_ (green, lower panels). Arrow points to the segment of the portal vessels used for measurements shown in **b**. **b**, Plot of AVP_FL_ intensity over time following its I.V. infusion (arrow). **c**, 2-photon image of SCN-OVLT portal vessels in another rat showing the presence of AVP_FL_ inside the vessels. **d**, Summary plots showing SON-OVLT portal vessel diameter (left) and blood flow velocity (right) at baseline and following the AVP_FL_ infusion. Vertical and horizontal arrows in A1 point rostrally and medially, respectively. Error bars represent ±SEM.

## DISCUSSION

The present study demonstrates unequivocally that blood flows from the SCN to the OVLT via a vascular portal pathway. Most importantly, blood flow in the SCN-OVLTp is regulated in a circadian manner. Importantly, AVP, a functionally relevant neuropeptide synthesized and secreted by the SCN, flows within the SCN-OVLTp. As noted in the Introduction, transplants of the copolymer encapsulated SCN that permit diffusion of neurosecretions but prevent establishment of neural connections confirm that the SCN produces sufficient diffusible neurosecretions to sustain circadian locomotor rhythms (11). The conclusion is supported by *in vitro* co-culture studies indicating that AVP, VIP and GRP from a wild-type SCN can drive rhythms in a target tissue of an SCN deficient mutant animal (15). Taken together, this small nucleus, comprised of ∼20,000 small neurons, generates chemical secretions of sufficient concentration to support circadian rhythms in nearby targets. Significantly, the present work points to the SCN-OVLTp as a route whereby such signals can travel.

Our findings are also consistent with prior work on the possible target site of a diffusible signal. The weight of evidence indicates that the target of diffusible SCN secretion lies nearby this nucleus. Thus, SCN transplants are successful in restoring rhythms when placed anywhere within the third ventricle (23) but not in remote sites. Calculations on the distance that diffusible SCN neurosecretions could travel suggest that the humoral signals have to act on a nearby target (24). The OVLT, lying at a distance of 385 µm from the SCN in mouse (2) and 730 µm in the rat meets the criterion. That the SCN and the OVLT constitute the source and target, respectively, within the SCN-OVLTp system is unequivocally determined by our *in vivo* blood flow measurements.

Importantly, we found that blood flow in this portal system is higher during nighttime compared to daytime, supporting the notion that blood flow in this system is amenable to modulation by rhythmic circadian signals. Our results showed that changes in blood flow occurred in the absence of changes in portal vessel diameter. Given that these are non-resistive vessels, these data suggest that mechanisms other than changes in the portal vessel diameter themselves contribute to blood flow regulation in this system.

Of immediate interest is the possible signal carried by the portal vasculature. We show that systemically infused fluorescently tagged AVP travels within the portal vessels. For several reasons, we focused on AVP although many other neurosecretions may course within the portal pathway. First, the OVLT has receptors for AVP (25). Furthermore, AVP is produced within the SCN with a daily rhythm, is a well-established SCN output signal (reviewed in (12, 25)) and can act in a diffusible manner (11-15, 23, 26). Rhythms in CSF AVP are lost in the absence of rhythms of a molecular clock controlling AVP in the SCN (27, 28). Finally, AVP plays an important role as an intercellular communication signal (29, 30), and is essential for SCN network synchrony and rhythmicity (9).

In summary, using a combination of anatomical and *in vivo* imaging techniques, we show that the SCN-OVLTp is a functional portal system that is present in both sexes, and carries circadian information from the SCN towards the OVLT. The SCN-OVLTp is a previously unknown new route and target for diffusible output signals from the brain clock in the SCN. The major anatomical features of the SCN-OVLTp pathway of the rat resembles that previously described in the mouse(2), and suggests that this pathway, like the pituitary portal system, is a universal feature in vertebrate brains.

## METHODS

### Animals

Male and female heterozygous transgenic eGFP-AVP Wistar rats (<u>Ueta et al.,</u> <u>2005</u>) (250–490 g, 8–16 weeks old) were used in all experiments (n = 12 and 16 females and males, respectively for *in vivo* studies; n= 2 and 2 females and males, respectively for iDISCO anatomical studies). Rats were housed in cages (2 per cage) under constant temperature (22 ± 2°C) and humidity (55 ± 5%) on a 12-h light cycle (lights on: 08:00– 20:00: [ZT0-ZT12]). In a subset of studies, rats were put on a reversed light cycle (lights on: 20:00–08:00). For anatomical studies, brains were shipped to Columbia University. All performed experiments were approved by the Georgia State University Animal Care and Use Committee (IACUC) and carried out in agreement with the IACUC guidelines. At all times, animals had ad libitum access to food and water and all efforts were made to minimize suffering and the numbers of animals used for the study.

### iDISCO clearing, light-sheet microscopy, image processing, vascular tracing

The iDISCO protocol used in this study is modified from(31). The tissue was labelled with anti-vasopressin (AVP) antibody to delineate the SCN, anti-smooth-muscle-actin (SMA) for arteries, and anti-type IV collagen for all blood vessels. Light sheet microscopic images were acquired with a LaVision Ultramicroscope II. Blood vessel tracing was done in Imaris and Vessulucida360. The SCN-specific labeling by AVP allows identification of the SCN without registration.

### *In vivo* 2-photon imaging of the SCN

A modified version of the transpharyngeal surgical approach to expose the ventral surface of the brain was used(32) in conjunction with in vivo 2-photon imaging as previously described(21). The exposed SCN (identified by the presence of eGFP-AVP neurons) was imaged under a 2-photon microscope excited with a Ti:Sapphire tuned at 860 and scanned with resonant galvanometers through a 16X (numerical aperture 0.8) water immersion objective or a 4X (numerical aperture 0.13) objective. Galvanometric scanning and resonant scanning (for imaging vessel dye loading and for RBC velocity) were controlled by PrairieView.

### *In vivo* imaging of the arterial and venous filling phases of the SCN vasculature

The artery-specific dye Alexa Fluor 633(22) (1–2 mg/kg injected I.V., femoral vein) 1.5 h before 2-photon imaging recordings was used to label and differentiate SCN arterioles from venules. Time-lapse imaging was performed using the resonant scanning mode at 30 Hz frame rate, 512 × 512 pixels during the intravenous administration of rhodamine or FITC 70 kDa dextrans (20 mg/ml, 200 nl/rat). Where noted, a fluorescently labeled form of arginine vasopressin (FAM-AVP, Anaspec) was infused in a similar way to the dextrans.

### *In vivo* quantification of blood flow rate and direction in the SCN-OVLT portal system

The SCN-OVLTp was identified as vessels originating from a dense SCN capillary network that run rostrally close to the midline towards the OVLT, and that were rhodamine/FITC 70 kDa dextran positive and Alexa Fluo 633 (an artery/arteriole specific dye)-negative. Blood velocity was calculated by monitoring the movement of red blood cells (RBC), which appeared as moving dark spots in fluorescently-labeled vessels). High-speed (500–1000 Hz, 5–10 s) resonant scanning images were obtained from a small region of interest (on average 150 × 25 pixels) drawn across the width of the vessel of interest. Velocity kymographs were constructed in ImageJ and used to determine directionality.

### Statistical analysis

Statistical analyses were performed using Prism 8 (GraphPad, CA, USA). Quantitative data were expressed as the MEAN ± SEM. No statistical methods were used to predetermine sample sizes, but sample sizes in the current study were similar to our previous reports(21, 33). Data were analyzed by paired or unpaired t test (two-sided in all cases). Chi-square tests were used to establish significant correlations. Data distribution was assumed to be normal, although this was not formally tested. Values were considered significantly different at p < 0.05. Statistical details of experiments can be found in the figure legends.

## SUPPLEMENTARY FIGURES

**Supplementary Figure 1-.**
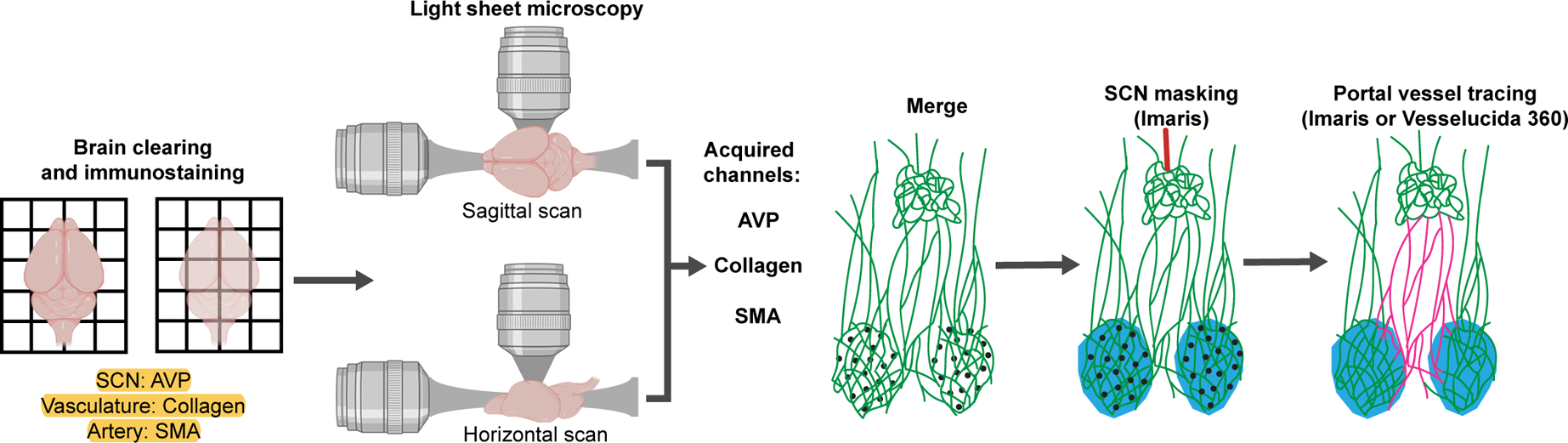
Experimental pipeline for iDISCO protocol. The cleared brains were labelled with AVP to localize the SCN, collagen to label the entire vasculature, and SMA to label the arteries. The samples were scanned either sagittally or horizontally. Because the vasculature of the OVLT is much denser than that of the SCN, a mask of SCN was created in Imaris to allow the visualization of SCN and the OVLT in the same scan. The collagen channel was used for blood vessel tracing. Created with BioRender (https://biorender.com/).

**Supplementary Figure 2-.**
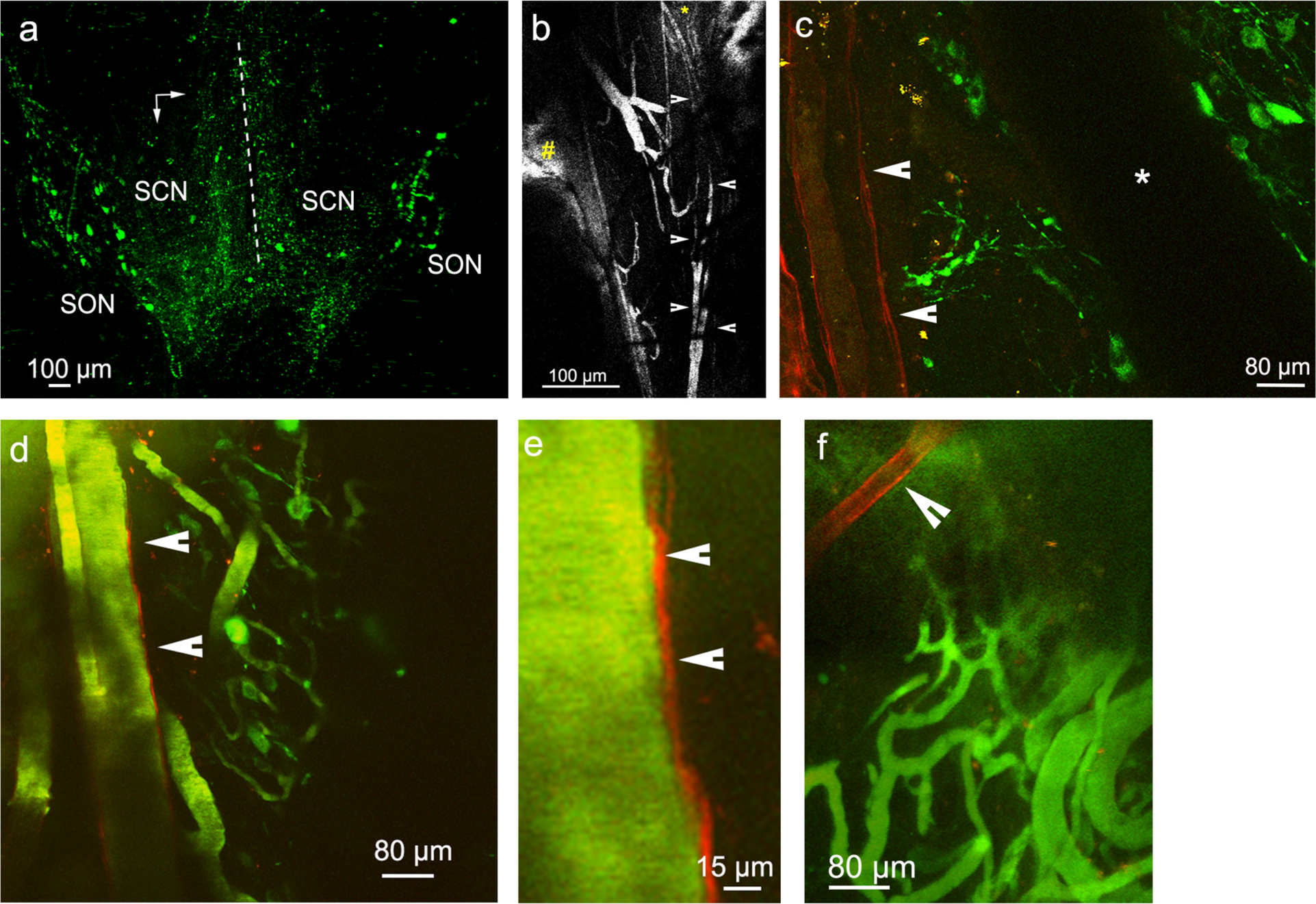
**a**, Composite of sequential medio-lateral images (4X objective) obtained in a rat in which the SON and SCN were exposed bilaterally. Dashed line indicates the apparent midline. **b**, Representative composite image at low magnification (4X objective) to better depict FITC 70 kDA dextran-<u>labelled</u> rostro-caudally running vessels emerging from the SCN along the midline (arrowheads). Note also the FITC 70 kDA dextran-<u>labelled</u> capillary vessels in the SCN (asterisk). The hashtag indicate the location of endogenous eGFP-AVP fluorescence in the SON. **c,** 2-photon image of the SON 1.5 h after I.V.. injection of the artery-specific dye Alexa Fluor 633. Note the Alexa Fluor 633 staining in a large penetrating arteriole (red, arrowheads) and the lack of staining in a large venule (asterisk). Endogenous eGFP-AVP fluorescent neurons are shown in green. **d**, 2-photon image of another SON showing Alexa Fluor 633 staining (arrowheads, red) in vessels subsequently stained with FITC 70 kDa dextran (I.V.,green). **e**, Same Alexa Fluor 633-stained vessel as in **d** shown at higher magnification. **f**, Same region as in ***Fig.4d*** showing at higher magnification the Alexa Fluor 633-stained arteriole (arrowhead) and FITC 70 kDa dextran stained capillaries and venules lacking Alexa Fluor 633 staining. Vertical and horizontal arrows in **a**, point rostrally and medially, respectively, and apply to all the images shown.

**Supplementary Video 1**: 2-photon imaging video of the SCN-OVLT portal vessels shown in Figure 8, depicting movement of RBCs moving in a caudal-to-rostral location. Vertical and horizontal arrows point rostrally and medially, respectively

## Author contributions

Y.Y., R.K.R., R.S. and J.E.S. designed the experiment and wrote the manuscript. Y.Y. and I.G conducted tissue clearing, acquired light-sheet microscope images and prepared figures, image processing, tracing. R.K.R conducted in vivo imaging experiments and R.K.R. helped in the preparation of figures.

## Acknowledgements

We thank Dr. Ferdinand Althammer (GSU) for general technical assistance.

## Data availability

Data associated with this study are present in the paper or in the Supplementary Information file. The raw data that support the findings of this study are available from the corresponding authors upon reasonable request. Source data are provided with this paper.

